# Ligand-Specified Signaling Efficacy Defined by Unique Transitions in G Protein Conformations

**DOI:** 10.64898/2026.07.23.740425

**Authors:** Xudong Wang, Wenkai Sun, Nathaniel Hays, Pangche Wang, Cheng Zhang, Yongbo Zhang, Libin Ye

## Abstract

Despite decades of research into the signaling efficacy of G protein-coupled receptors (GPCRs), including the resolution of numerous receptor structures in complex with nucleotide-free G proteins^1,2^, mini-G proteins^3,4^, and G protein carboxyl-terminal peptides^5–7^, our understanding of the underlying mechanisms remains elusive, because a fluidic, dynamic view of the conformational transitions during GPCR activation is still missing. Benefiting from its supersensitization to microenvironmental changes, recent advances in ^19^F-quantitative NMR (^19^F-qNMR)^8^ enable the dissection and quantification of the conformational energy landscape and dynamics of receptors and their signaling partners, offering new perspectives for deciphering the signaling machinery through a novel lens. Herein, we first established a 3-state model of Gα_s_ conformational landscape and then quantitatively investigated the conformational transitions and dynamics of Gα_s_ proteins in response to different ligand-bound receptors. Our research demonstrated that Gα_s_ substates are not only differentially populated in a ligand-dependent manner but also undergo distinct conformational-cycling kinetics across sub-states in response to different ligand bindings. These discoveries enable us to propose a novel GPCR model that integrates quantitative G protein conformational dynamics at the atomic level to explain GPCR pharmacological efficacy, emphasizing that unique equilibrium and transitions of G protein conformational states in response to ligands are critical determinants of signaling specificity.

## Main

The development of cryo-EM technologies has significantly advanced our ability to determine the static structures of G protein-coupled receptors (GPCRs) in complex with transducers such as heterotrimeric G proteins, GRKs, and β-arrestin, yielding abundant structural data^1,9–18^. However, our understanding of the dynamic, fluidic conformational transitions and their impacts on signaling outputs during GPCR activation remains elusive due to the difficulty of studying these aspects. This is particularly evident, as all resolved GPCR-G protein complex structures represent the fully activated state until recently, a nearly-activated intermediate GPCR-mini-G complex structure was resolved through the conformational transition blockage strategy^3,19^. Despite this progress, our understanding of GPCR signaling still largely relies on dotted structural snapshots, leaving a gap in connecting dynamic transitions to functional specificity. In our previous study, we characterized the conformational and dynamic behavior of the classic GPCR, the adenosine A2A receptor (A_2A_R), in response to ligand binding using ^19^F quantitative NMR (^19^F-qNMR)^3,8,19,20^. Nonetheless, our understanding of how its cognate G proteins respond to changes in receptor conformation and dynamics elicited by ligand binding remains elusive^21^. Here, we aimed to investigate the conformational landscape of G proteins at the quantitative level and examine their responses to distinct ligand-bound receptors. Our goal is to achieve a mechanistic understanding of how Gα_s_ conformational transitions among states and their rates contribute to signaling effectiveness, thereby expanding our view of GPCR signaling beyond static images of current GPCR-G protein complexes.

Recent studies indicate that intermediate GPCR-G protein complexes can conduct a rate-limited nucleotide exchange, either independent of or dependent on transitioning to the fully activated complex^3,22–24^. We thus hypothesize that both the populations and the dynamics of these signaling GPCR-G protein complexes, including fully activated and intermediate complexes, contribute to signaling efficacy. Accordingly, we aim first to delineate the conformational landscape of the G protein and then to quantify both the subpopulations and the dynamics of these substates in response to different ligand-bound receptors. To achieve this, we utilized a highly sensitive tri-fluorinated tag, 2-bromo-N-(4-(trifluoromethyl)phenyl)acetamide (BTFMA)^25–27^, to probe the conformational behavior of the Gα_s_ subunit. Research has indicated that one of the main features of receptor- activated G protein is the insertion of its Cα5 helix into the G protein-binding cavity. ^5,9,28^ (Figs. 1A and 1B). Thus, we aimed to depict this dynamic motion process using ^19^F-qNMR in response to different ligands (full, partial, and inverse agonists). To prevent the nonspecific ^19^F labeling, a cysteine-minimized construct^29^ (Gα_s_-Δ6: C3S-C200T-C237S- C359I-C365A-C379V) was utilized, and single cysteine substitutions along the Cα5 helix for site-specific fluorine tagging were introduced (Fig.1C). To minimize the structure- function perturbation caused by mutation and ^19^F-tag labeling, all variants were expressed, labeled with BTFMA (Fig.1E), and their activities were evaluated relative to the wild-type protein. Unexpectedly, the original Gα_s_-Δ6 construct exhibited a significantly reduced GTP hydrolysis activity compared with the wild-type (Extended Data Fig. 1), whereas the introduction of a cysteine at position R380 (Gα_s_-Δ6_R380C) significantly restored the GTP turnover functionality and reached a comparable capacity to the WT construct in the 2h hydrolysis experiments (Fig. 1D and Extended Data Fig.2). BODIPY-FL-GDP binding assessments (Figs. 2A, 2D, and 2E) indicated that the GDP association and dissociation processes of the Gα_s_-Δ6_R380C mutant were much slower than the WT, in which T_peak_ (time to peak binding) shifted from 23.5 min to 29.0 min while T_eq_ (time to equilibrium) shifted from 40.75 min to 46.0 min in apo sample. However, within the Gα_s_-Δ6_R380C set, the GDP-binding pattern and nucleotide exchange in response to various ligands are consistent with those of the WT (Figs. 2F and 2G). That implies that the Gα_s_-Δ6_R380C construct can recapitulate the signaling process of the wild type, albeit at a slower rate. Notably, the prolonged GDP association and dissociation process may result from both the mutations of the C365A in the conserved guanine-interacting TCAT/V motif (Fig. 2C) and the R380C on the Cα5 domain. Although R380C is not a key residue that interacts with the receptor, it weakens receptor-G protein interaction^4^, resulting in decreased opening of the AHD from the Ras domain of the G protein. This was reflected in prolonged GDP association and dissociation processes (increased values of T_peak_ and T_eq_), along with a relatively lower initial rate of GTPγS replacing BODIPY-FL-GDP. This observation indicated that GDP binding to the G protein is influenced not only by GDP-binding pocket residues, such as C365^30–33^, but also by the receptor’s modulatory capacity on the G protein, which plays a critical role in conveying G protein activation. Given the same trend in the Gα_s_-Δ6_R380C response to the ligands for GDP binding and nucleotide exchange despite affinity difference, we concluded that the insertion motion process of Cα5 into the intracellular G protein-binding pocket can be monitored via the 19F-probe at the R380C of the Gα_s_ (Figs. 1A and 1B). Successful conjugation of BTFMA to the Gα_s_-Δ6_R380C was evaluated by LC-TOF mass spectroscopy (Extended Data Fig.3 and Extended Data Table 1). ^19^F-NMR spectra further indicated that a non-cysteine residue in the Gα_s_-Δ6 is accessible for labeling, where the ^19^F-tag can specifically be attached to the construct of Gα_s_-Δ6_R380C (Extended Data Fig.4).

**Fig. 1.**
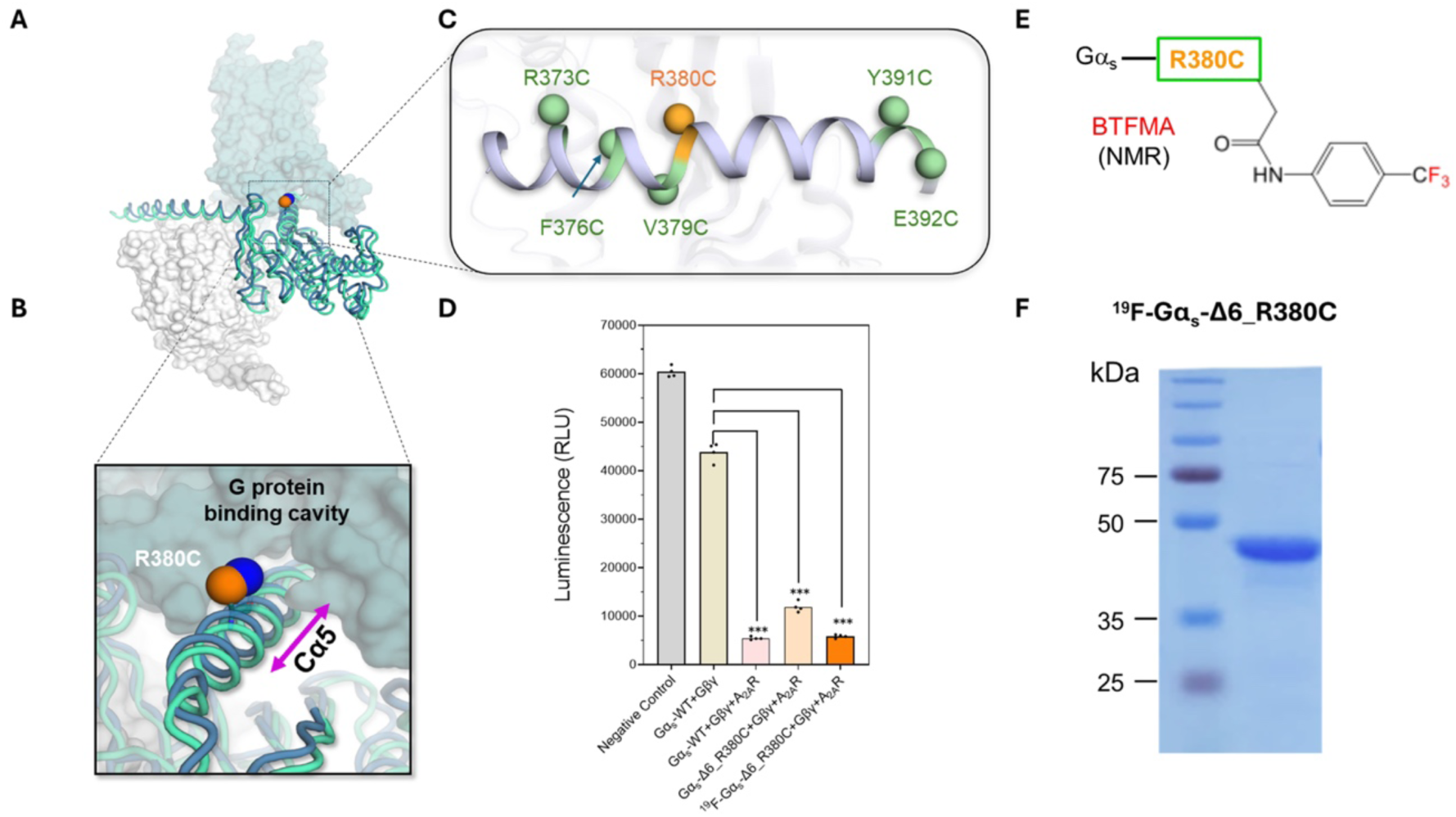
The strategy of creating Gα_s_-Δ6_R380C for studying the conformational transitions and dynamics when heterotrimeric G proteins engage the receptor. **A**. The overview of GPCR-G protein complex and the location of residue R380C in the interfacial cavity. **B.** Proposed the motion of the Cα5 (and residue R380C) helix as the G protein inserts and retracts from the G protein binding pocket. **C**. Screened sites on the Cα5 helix for ^19^F-labeling. **D**. GTP conversion capacity in 2 hours of ^19^F-labeled constructs for Gαs-Δ6 mutants and wild-type. **E**. Structure of protein-attached ^19^F-label for NMR study — BTFMA: 2-bromo-N-(4- (trifluoromethyl)phenyl)acetamide. **F**. SDS-PAGE of the expressed Gα_s_-Δ6_R380C. NF: non-^19^F labeling; ^19^F: BTFMA labeling.

**Fig. 2.**
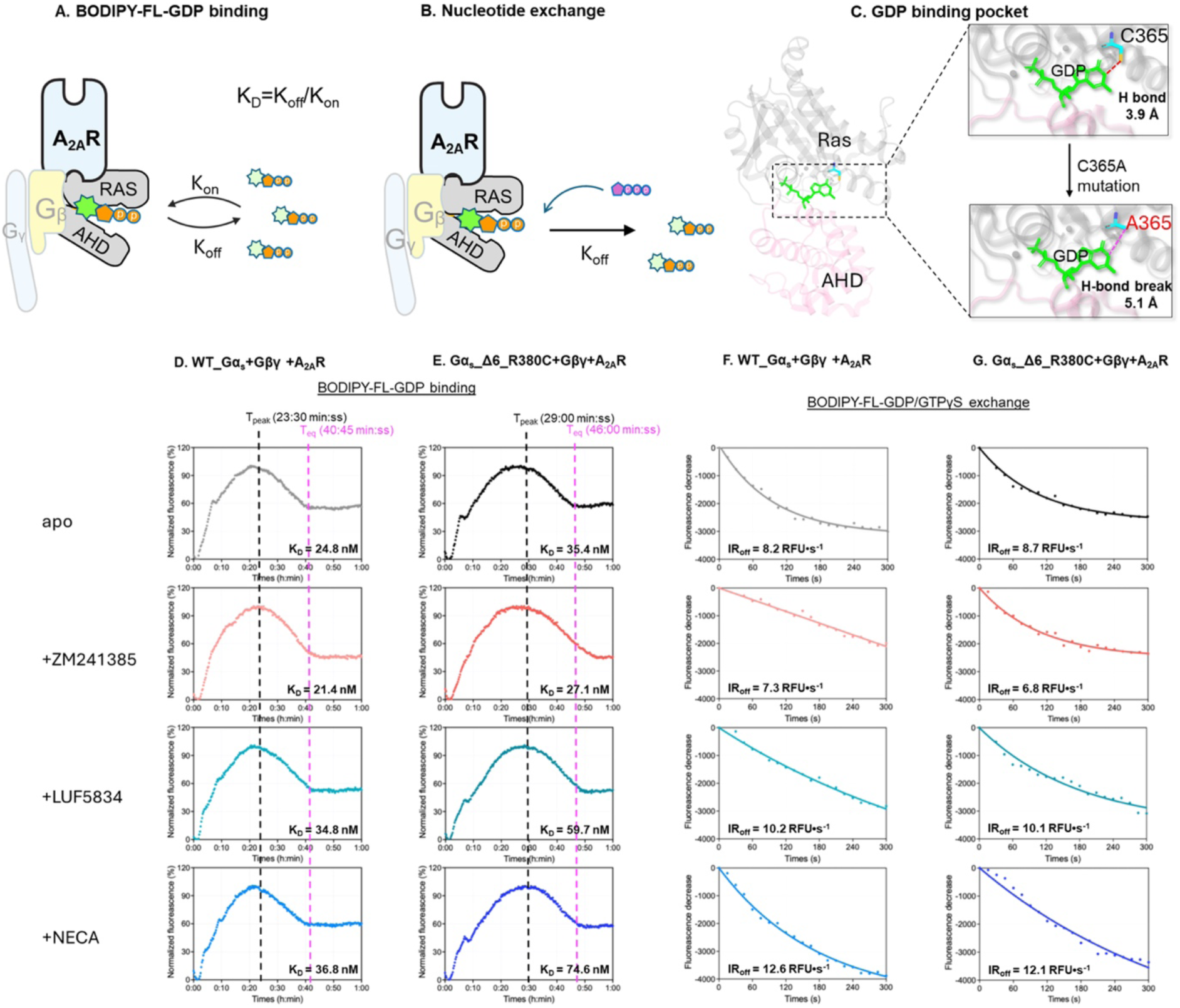
Mutant Gα_s_-Δ6_R380C shows the same trend but relatively slower rates of nucleotide binding and exchange. **A**. A cartoon model of BODIPY-FL-GDP binding assay. **B**. A cartoon model of nucleotide exchange assay by replacing BODIPY-FL-GDP with GTP. **C**. The depiction of the nucleotide-binding pocket in the WT and mutant with C365A in TCAT/V motif. **D**. BODIPY-FL-GDP binding assays for WT in response to inverse agonist (ZM241385), partial agonist (LUF5834), and full agonist (NECA), along with the apo sample. **E**. BODIPY-FL-GDP binding assays for Gα_s_-Δ6_R380C in response to inverse agonist (ZM241385), partial agonist (LUF5834), and full agonist (NECA), along with the apo sample. **F**. Nucleotide exchange for WT in response to inverse agonist (ZM241385), partial agonist (LUF5834), and full agonist (NECA), along with the apo sample. **G**. Nucleotide exchange for Gα_s_-Δ6_R380C in response to inverse agonist (ZM241385), partial agonist (LUF5834), and full agonist (NECA), along with the apo sample. Of note: IR_off_ represents the initial replacement rate of BODIPY-FL-GDP by GTPγS.

We acquired 1D ¹⁹F NMR spectra of Gα_s_ in complex with Gβγ and the A_2A_ receptors, pre- saturated with the inverse agonist ZM241385, the partial agonist LUF5834, the full agonist NECA, and the unliganded (apo) state. Three distinct conformational components of Gα_s_ were observed and orchestrated with the ligands. Since these components concerted with ligand bindings, we designated them as the free/pre-coupled state (P1), the partially engaged state (P2), and the fully engaged state (P3) (Fig. 3A-C, and Extended Data Fig.5). Of note, the G protein structures from PDB IDs: 6EG8 (red, inactive)^5^, 9EE8 (teal, intermediate)^3^, and 6GDG (blue, fully active)^34^ were used for modeling. The chemical shift rebalances from P1 (high magnetic field, the most solvent-exposed) to P3 (low magnetic field, the least solvent-exposed), suggesting that the R380C residue becomes more hydrophobic/electronically shielded as it approaches the binding cavity. Furthermore, we quantified the subpopulation of each state (Figs. 3D-E). When the G proteins engaged the receptors that were saturated with the inverse agonist ZM241385, the P1 state of the Gα_s_ was predominantly populated. In contrast, the P3 fraction was further populated in the presence of a full agonist NECA, whereas the P2 fraction was more prominent in the case of LUF-saturated receptors. It is essential to note that, in all cases, we initially set the G protein-to-receptor ratio at 1:5, anticipating that all G proteins bind not only to the receptor but primarily to a single receptor conformation. However, this was not the case, indicating that G protein binding to one receptor conformational state leads to a reciprocal, balanced transition to the other state, rather than a single binding event to a specific receptor state.

**Fig. 3.**
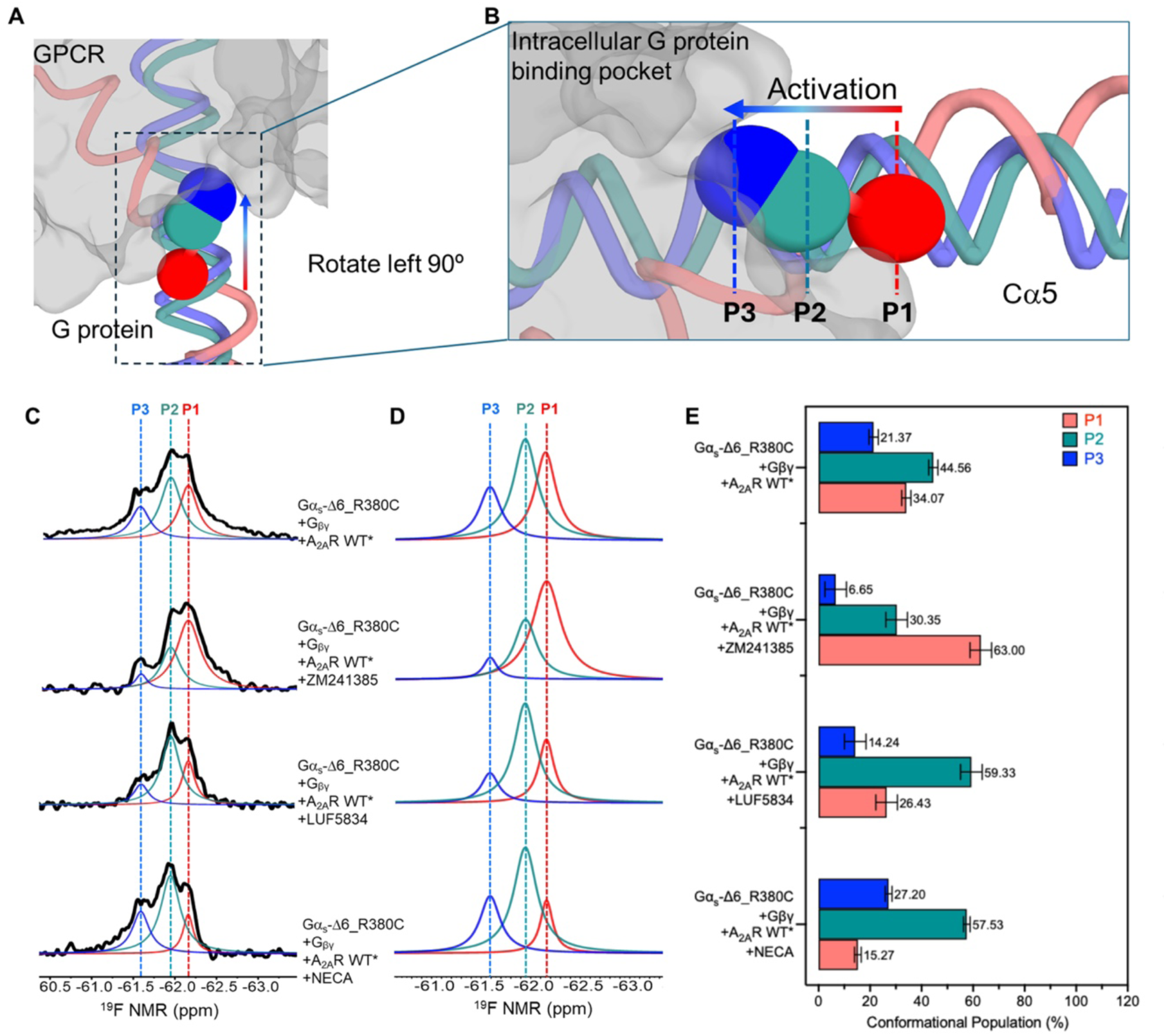
Conformational transitions of the Cα5 helix from Gα_s_ in response to different ligand-bound A_2A_R receptors, profiled by ^19^F-qNMR through labeling at R380C. **A**. The side-view of the Cα5 motif engages the intracellular G protein binding pocket of the GPCR. The colors of red, turquoise, and blue represent three states of the Cα5 in the G protein activation process. The PDBs: 6EG8 (red, inactive), 9EE8 (teal, intermediate), and 6GDG (blue, fully active) were used for modeling. **B**. Highlighted portions of the Cα5 helix and its corresponding peaks in ^19^F-qNMR spectra, shown in panel **C**. **C**. Conformational profiles of the Cα5 helix probed by ^19^F-qNMR in response to inverse agonist (ZM241385), partial agonist (LUF5834), and full agonist (NECA)-bound receptor. **D**. Deconvolution of conformational states of the Cα5 helix probed by ^19^F-qNMR. **E**. The populations of each conformational state of the Cα5 helix in response to different ligand bindings.

Our previous study indicated that the transition from the inactive state to the active (-like) states of the receptor took 0.7±0.1 seconds upon full agonist binding, whereas it took 2.7±1.3 seconds with partial agonist binding^8^. Due to the substantial overlap between the fully active and active-like states in the receptor’s conformational profile, we were unable to precisely determine the transition time from the inactive to the fully active state^8^. However, a prolonged transition time can be expected due to an extended transition trajectory from the inactive to the fully active state, compared to a comprised active(-like) conformational ensemble. With this in mind, we wondered whether G proteins transition to the binding pocket at different rates in response to different ligands. The rate of Gαβγ conformational transitions is expected to correlate with signaling efficacy; accordingly, defining the differences in these transition processes is essential for understanding variations in signaling output. However, this type of information is lacking in the field for several reasons, including the challenge of large amounts of receptor preparation for NMR studies^35,36^, and macromolecular ¹⁹F-NMR is limited by inherently low signal-to-noise ratios despite its pronounced sensitivity to local environmental changes ^37,38^.

Building on this concept and using the G conformational landscape established above, we further employed relaxation experiments—¹⁹F-chemical exchange saturation transfer decay (¹⁹F-CEST)—to assess the transition rates from P3 to P1. The working principle of ^19^F-CEST was illustrated in Extended Data Fig. 6, where a radiofrequency (RF) pulse saturating one of the relevant conformational states will affect the second state’s behavior due to chemical exchange between these two components. Notably, the significant spectral overlap between P1 and P2 prevents selective saturation of either state, thus hindering direct measurement of their individual transitions to P3. Instead, we designed CEST experiments (Figs. 4A-D) to examine the lifetime of the P3 component and its chemical exchange with the P2 component by saturating P3 and tracking the decay of P2. The chemical exchange rate between P3 and P2 can be derived from the P2 relaxation with mixing times ranging from 0 to 1.2 seconds. The exchange rate from P3 to P2 when G protein engages the NECA-bound receptor was found to be 10.39 ± 3.52 s⁻¹ (Fig. 4B). Conversely, the conformational exchange rate of Gα_s_ in the LUF5834-bound receptor decreased to 6.26 ± 3.96 s⁻¹ (Fig. 4C). Prior studies indicated that receptors bound to inverse agonists continue to sample multiple conformational states^8,39,40^, and similar behavior was observed in this study, suggesting no ligand can completely restrict the G-protein conformational ensemble to a single state. When G proteins engaged receptors saturated with the inverse agonist ZM241385, the population of fully active GPCR-G protein complexes significantly dropped, accompanied by a population increase in the P1 state and a substantial reduction in the exchange rate to 1.09 ± 1.08 s⁻¹ (Fig. 4D). The lifetime of P3 was calculated using the formula τ = 1/K, referencing the decay of P2. It was 0.10 s, 0.16 s, and 0.90 s when G protein engaged full, partial, and inverse agonist-bound receptors, respectively (Figs. 4B-D).

**Fig. 4.**
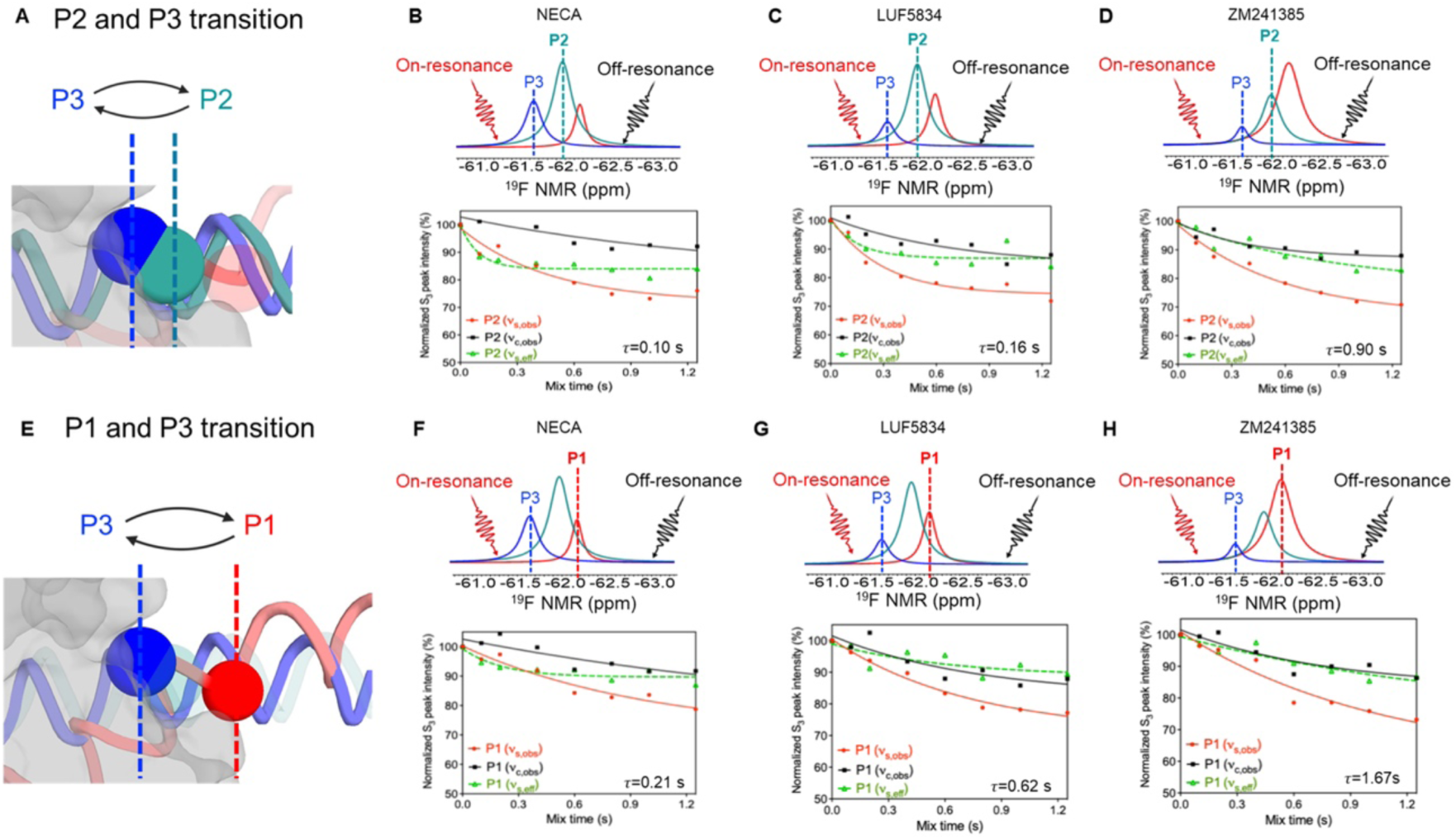
G protein dynamics during receptor activation measured by ^19^F-qNMR CEST decay. **A**. The cartoon representation of the dynamic motion between P2 and P3 states using ^19^F-labeled R380C as a probe. **B.** Conformational exchange rate between P3 and P2 of G proteins when bound to a full agonist-NECA saturated receptor, and the lifetime of the G protein-bound state P3 as measured by P2 decay. **C.** Conformational exchange rate between P3 and P2 of G proteins when bound to a partial agonist-LUF saturated receptor, and the lifetime of the G protein-bound state P3 as measured by P2 decay. **D.** Conformational exchange rate between P3 and P2 of G proteins when bound to an inverse agonist- ZM241385 saturated receptor, and the lifetime of the G protein-bound state P3 as measured by P2 decay**. E.** The cartoon representation of the dynamic motion between P1 and P3 states using 19F-labeled R380C as a probe. **F.** Conformational exchange rate between P3 and P1 of G proteins when bound to a full agonist-NECA saturated receptor, and the lifetime of the G protein-bound state P3 as measured by P1 decay**. G.** Conformational exchange rate between P3 and P1 of G proteins when bound to a partial agonist-LUF saturated receptor, and the lifetime of the G protein-bound state P3 as measured by P1 decay**. H.** Conformational exchange rate between P3 and P1 of G proteins when bound to an inverse agonist- ZM241385 saturated receptor, and the lifetime of the G protein-bound state P3 as measured by P1 decay.

Likewise, we examined the conformational transitions between the conformers P3 and P1 (Figs. 4E-H). The lifetimes of the P3 in response to ligands NECA, LUF5834, and ZM241385 could also be measured based on the P1 decay, resulting in 0.21 s, 0.62 s, and 1.67 s (Figs. 4F-H). It is worth noting that the different lifetimes of P3 obtained from the decays of P2 and P1 were due to the two-state model applied for these decay measurements. However, this also offers an easy way to calculate the exchange time from P1 to P2 by subtracting the P3 lifetime in the P3-P2 exchange system from that in the P3-P1 exchange system. Our research clearly shows that the complex takes longer to move from the pre-coupled state to the partially activated state, and then to the fully activated state, in systems bound to partial agonists than in those bound to full agonists.

## Conclusion

Although pharmacological studies have established that full agonist binding elicits maximal signaling whereas partial agonist binding produces submaximal signaling, the mechanistic basis for this difference remains elusive^3^, particularly in terms of how the time kinetics of different state transitions contribute to signaling levels. This study examined G-protein dynamics by quantifying conformational substates and their transitions in response to receptors bound to ligands of differing efficacy, providing mechanistic insight into GPCR-G-protein signaling ^8^. Here, a mechanistic model is proposed (Fig. 5) that describes how ligand binding reshapes the subpopulations and transition dynamics of Gα_s_ conformations, thereby linking these features to pharmacological efficacy. When Gα_s_ binds to the receptor with a full agonist, the fully engaged state, P3, is the most prevalent. Conversely, in the presence of a partial agonist, Gα_s_ predominantly occupies the P2 state. In both cases, enrichment of either P3 or P2 occurs at the expense of the remaining substates. This indicates that the overall signaling efficiency results from a combination of GPCR-P3 and GPCR-P2 complexes, which are combinatorially determined by their state populations and transition rates in response to different ligands. In addition, each single cycling trajectory of the G protein is much shorter than that of the receptor, implying that multi-cycling G protein binding events could occur within each receptor activation trajectory, another factor contributing to signaling efficacy. These findings offer a quantitative, dynamic framework for interpreting GPCR signaling, showing that unique, ligand-specified G protein conformational transitions and dynamics contribute to functional specificity.

**Fig. 5.**
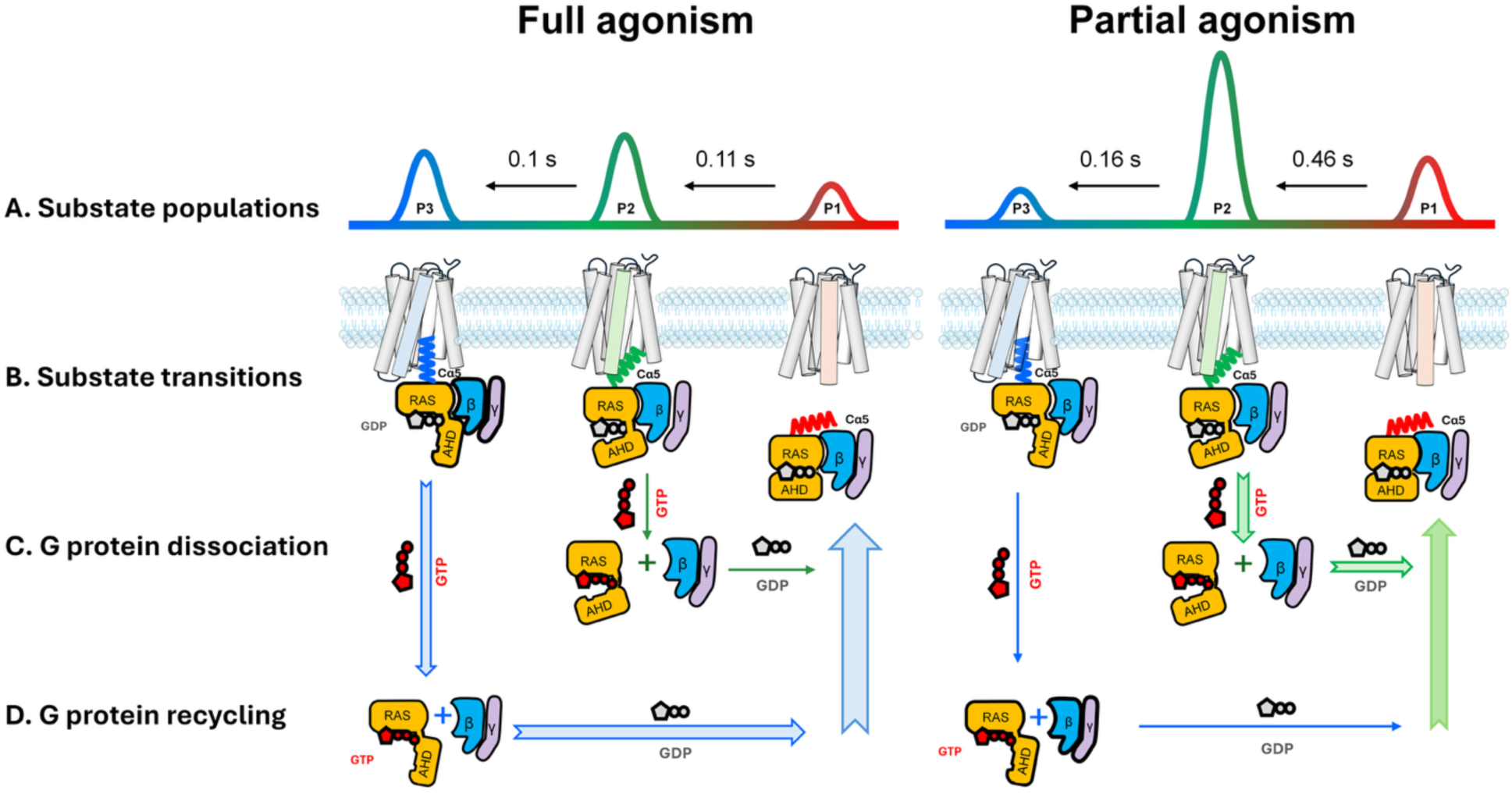
A proposed mechanism for GPCR signaling efficacy involving quantitative G protein conformational changes and dynamics. This model demonstrates that signaling efficiency depends not only on the populations of each substate but also on the rates of dynamic transitions between them in response to different ligands. Note: the thickness of the arrow represents the strength of the signaling level.

## Materials and Methods

### Receptor Expression and Purification

The full-length human *ADORA2A* gene (A_2A_R), derived from the pPIC9K_ADORA2AR construct generously provided by Prof. Takuya Kobayashi (Kyoto University), was used to generate the A_2A_R_316 variant incorporating an N-terminal FLAG tag and a C-terminal His-tag, as described previously.

Plasmids were linearized with PmeI-HF (New England Biolabs) and electro-transformed into *Pichia pastoris* SMD1163 (Invitrogen; Δhis4Δpep4Δprb1) using a Gene Pulser II (Bio-Rad). High-copy clones were selected on YPD plates containing G418 at 1, 4, and 6 mg/mL. Colonies from 6 mg/mL plates were screened via immunoblotting with anti-FLAG and anti-poly-His antibodies. Antibodies were validated according to the manufacturer’s datasheets.

Transformants expressing wild-type were pre-cultured on YPD agar with 0.1 mg/mL G418. A single colony was inoculated into 4 mL of YPD medium and incubated at 30°C for 12 h. It was then transferred into 200 mL of BMGY medium and cultured for 30 h at 30 °C. Cultures were scaled up to 1 L of BMMY medium containing 3% DMSO and 10 mM theophylline and induced with 0.5% methanol at 20 °C. Methanol was replenished every 12 h for 60 h before harvest. Cells were pelleted (4000 × g, 20 min), washed with 50 mM HEPES (pH 7.4), 10% glycerol, and lysed in breaking buffer (50 mM HEPES, pH 7.4, 100 mM NaCl, 2.5 mM EDTA, 10% glycerol) using a Microfluidizer (3 passes, 20,000 psi). Membranes were isolated by ultracentrifugation (100,000 × g, 2 h), resuspended in lysis buffer (50 mM HEPES pH 7.4, 100 mM NaCl, 1% MNG-3, 0.2% CHS), and solubilized at 4 °C for at least 2 h. Solubilized membranes were incubated with Talon resin (Clontech) for 2 h at 4 °C, washed with 50 mM HEPES pH 7.4, 100 mM NaCl, 0.02% MNG-3, and 0.01% CHS, and then eluted with 250 mM imidazole in the same buffer. Aliquots were flash-frozen and stored at −80 °C.

### Gα_s_ Expression, Purification, and ^19^F Labeling

The full-length human wild-type GNAS gene fused to MBP with an N-terminal 10×His tag was synthesized by Gene Universal. An engineered GS flexible linker along with a TEV protease cleavage site was inserted between MBP and Gαs. Overlap-extension PCR (OE-PCR) using primers Gαs-F-NcoI and Gαs-R-KpnI was used to generate the MBP- Gαs fusion, which was then digested with NcoI and KpnI and ligated into the pBAD backbone under the control of the arabinose-inducible araBAD promoter.A series of cysteine-light and site-specific mutants were created from this construct. The cysteine- light background (Gαs-Δ6) was generated by introducing mutations C3S, C200T, C237S, C359I, C365A, and C379V. Additional site-specific substitutions, including R373C, F376C, R380C, V379C, Y391C, and E392C, were introduced individually or in combination to enable site-specific ^19^F labeling (All primers and plasmids summarized in Extended Data Tables 2 and 3). Mutagenesis was carried out using the QuikChange Lightning Multi Site- Directed Mutagenesis Kit (Agilent Technologies). All constructs were confirmed by Sanger sequencing (Eurofins Genomics, USA).

For protein expression, the pBAD-MBP-Gα_s_ variants were transformed into *E. coli* DH10B competent cells and selected on LB agar containing 50 µg/mL ampicillin. Overnight cultures were inoculated into 2×YT medium with antibiotics and grown at 37 °C until the optical density at 600 nm (OD_600)_ reached ∼0.6. The cultures were then cooled to 20 °C, induced with 0.1% L-arabinose, and incubated at 18 °C for 24 h before harvesting. Cells were harvested (8000 rpm, 5 min, 4 °C), washed with 50 mM HEPES pH 7.4, 100 mM NaCl, and lysed in buffer containing 50 mM HEPES pH 7.4, 100 mM NaCl, 0.02% MNG- 3, 0.001% CHS, 1 mM MgCl_2_, 100 µM GDP, and 20 mM imidazole. Lysate was clarified by centrifugation. Clarified lysate was incubated with Talon resin (2 mL/L culture) for 2 h at 4 °C. Resin was labeled with 10-20 molar excess BTFMA (Apollo Scientific) for 24 h at 4 °C. Bound protein was eluted using 300 mM imidazole. TEV protease (1:50–1:100 w/w) was added following dialysis into cleavage buffer (50 mM HEPES pH 7.4, 100 mM NaCl, 0.02% MNG-3, 0.001% CHS, 1 mM MgCl_2_, 100 µM GDP, 0.5 mM EDTA, 1 mM DTT) and incubated overnight at 4 °C. Cleaved protein was collected from Talon resin flow-through and buffer-exchanged into storage buffer (50 mM HEPES pH 7.4, 100 mM NaCl, 0.02% MNG-3, 0.001% CHS, 1 mM MgCl_2_, 100 µM GDP). Aliquots were flash-frozen and stored at -80 °C.

### Gβγ Purification from Insect Cells

Gβγ subunits were expressed in insect cells and purified as follows: Cell pellets from 1 L of culture were washed with 150 mL lysis buffer (10 mM Tris-HCl pH 7.5, 100 µM MgCl₂, 5 mM β-mercaptoethanol) and centrifuged at 12,000 rpm for 15 min. The resulting membrane pellets were resuspended in 80 mL solubilization buffer (20 mM HEPES pH 7.4, 1% sodium cholate, 100 mM NaCl, 0.2% MNG, 5 mM MgCl₂, 2 µL calf intestinal phosphatase (CIP), 5 mM β-mercaptoethanol, and protease inhibitor cocktail).

Cell suspensions were mixed with glass beads and agitated at maximum speed for 2 h at 4 °C, then centrifuged at 12,000 rpm to remove debris. The supernatant was incubated with 5 mL of Talon resin for 2 hrs at 4°C with gentle shaking. Resin was collected by brief centrifugation (700-2000 rpm, 1 min), transferred to a gravity-flow column, and washed with 20 mL solubilization buffer followed by washing buffer (20 mM HEPES pH 7.4, 100 mM NaCl, 0.05% MNG-3, 1 mM MgCl_2_, 2 µL CIP, 5 mM β-mercaptoethanol, protease inhibitor cocktail). Bound proteins were eluted with a washing buffer supplemented with 300 mM imidazole. Eluates were concentrated and dialyzed overnight at 4 °C against at least 100-fold volume of washing buffer. Final purification was performed using ion exchange FPLC with a low salt buffer A (20 mM HEPES pH 7.4, 40 mM NaCl, 0.05% DDM, 1 mM MgCl_2_, 100 µM DTT) and a high salt buffer B (buffer A supplemented with NaCl to 1 M).

### 19F NMR and Chemical Exchange Saturation Transfer Experiments

NMR samples (280–300 µL) were prepared in 50 mM HEPES buffer (pH 7.4) containing 100 mM NaCl, 0.02% MNG-3, 0.002–0.01% CHS, and 10% D₂O. Samples contained 20– 50 µM A₂_A_R and, where specified, 25 µM bendroflumethiazide was added as a chemical shift reference at −59.05 ppm. For ligand-binding experiments, 1 mM ligand was added, and samples were incubated at room temperature for 20 min prior to measurements. All samples were loaded into Shigemi tubes. ^19^F NMR spectra were acquired at 20 °C on a 600 MHz Varian Inova spectrometer equipped with a dedicated ^19^F probe, using a 16 µs 90° excitation pulse, 200 ms acquisition time, 15 kHz spectral width, and a 1 s recycle delay. Typically, 15,000-50,000 scans were collected. Data were processed using VnmrJ 4.2 with zero filling and exponential apodization equivalent to 15 Hz line broadening, and analyzed with MestReNova 14.2.

To investigate slow exchange between resolved states, ^19^F chemical exchange saturation transfer NMR experiments were performed, in which a series of continuous wave (cw) irradiation pulses were applied at both an on-resonance frequency (Ѵ_s_) and off-resonance frequency (Ѵ_c_), to assess chemical exchange during steady state saturation and off-resonant saturation effects, as shown in Extended Data Fig. 6. Upon saturating the resonance associated with state B, the ideal magnetization response of A maybe described by the following formula^41^:

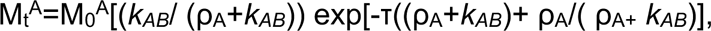

assuming off-resonant effects are accounted for. Note that both the exchange rate constants, *k_AB_*, and the longitudinal relaxation rate of spin A, ρ_A,_ can in principle be calculated from a fit of the above equation to the experimental data. Accordingly, the lifetime τ_A_ can be calculated from τ=1/*k_AB_*. All fits were performed using Gnuplot (www.gnuplot.info).

### GTPase Hydrolysis Assay

The GTPase hydrolysis assay was performed using a modified GTPase-Glo™ assay (Promega). Reactions for comparison of Gα_s_ variants were performed by mixing 300 nM Gα_s_-WT or the indicated Gα_s_ mutants with 300 nM Gβγ and 300 nM A_2A_R WT* in a final volume of 10 μL containing 50 mM HEPES (pH 7.4), 100 mM NaCl, 0.002% (w/v) CHS, and 0.02% (w/v) MNG-3. After incubation for 30 min at room temperature, 10 μL of 2× GTP-GAP solution containing 10 μM GTP, 1 mM DTT, and the cognate GAP was added, followed by incubation for 120 min at room temperature. Subsequently, 20 μL of reconstituted GTPase-Glo™ reagent containing 5 μM ADP was added, and the mixture was incubated for 30 min with shaking. Luminescence was measured after the addition of 40 μL detection reagent and incubation for 10 min using a VANTAstar F plate reader. RLU values were normalized to buffer controls and analyzed using GraphPad Prism 10. Each experiment was independently repeated three times.

### BODIPY-FL-GDP Nucleotide-Binding and Exchange Assays

The nucleotide-binding assay was performed using BODIPY-FL-GDP (Invitrogen™), supplied as a mixture of two isomers with the fluorophore attached at either the 2′ or 3′ position of the ribose ring. All measurements were conducted in assay buffer containing 20 mM HEPES (pH 7.4), 1 mM EDTA, 10 mM MgCl₂, and 0.01% (w/v) MNG. Receptor– G protein mixtures were assembled by combining purified Gα_s_, Gβγ, and A_2A_R WT* directly in assay buffer to yield 20 µM G protein (Gα_s_ with Gβγ) and 20 µM receptor in a 10 µL pre-incubation volume. Ligands (NECA, ZM241385, or LUF5834) were added at 800 µM, and equivalent volumes of DMSO were added to apo controls. Pre-incubation was performed for 20 min at room temperature.

For nucleotide-binding measurements, 90 µL assay buffer containing 100 nM BODIPY-FL-GDP was dispensed into 96-well half-area microplates, and reactions were initiated by the addition of 10 µL pre-equilibrated protein mixtures followed by rapid mixing. Fluorescence was recorded at room temperature using a VANTAstar F plate reader (excitation 475 nm; emission 528 nm). A no-protein control trace recorded under identical conditions was subtracted from all experimental traces to correct for the slow baseline decay of BODIPY-FL-GDP fluorescence. For cross-condition comparison, background-corrected fluorescence traces were normalized by defining the mean fluorescence during the first 70 s in the absence of protein as 0% and the maximum fluorescence observed in each individual trace as 100%. Normalized kinetic data were fitted using an association–then–dissociation model in GraphPad Prism.

Nucleotide exchange reactions were initiated directly from the nucleotide-binding assay following the 60 min incubation with BODIPY-FL-GDP by the addition of unlabeled GTPγS (Invitrogen™) at a 10-fold molar excess relative to BODIPY-FL-GDP. Fluorescence was monitored continuously at room temperature for up to 60 min. Baseline-corrected fluorescence traces were adjusted by subtracting the fluorescence value at 0 s for each individual trace, such that the initial fluorescence was set to 0 RFU and subsequent changes were expressed relative to this starting value. The initial 5 min of each decay trace were fitted to a one-phase exponential decay model with Y0=0 in GraphPad Prism, and initial rates were calculated from the mean fluorescence change over the first 4 min.

### LC–MS Analysis

Three Gαs protein samples were analyzed: the wild type (WT, Arg380), the R380C mutant, and the R380C mutant pre-labeled with the BTFMA probe during preparation. Prior to denaturation, residual free thiols were capped with iodoacetamide (IAM, Thermo Scientific Chemicals) at 25 mM, adjusted to pH ∼8, and incubated for 20-30 min at room temperature in the dark. Proteins were then denatured by addition of urea to 8 M and reduced with tris(2-carboxyethyl)phosphine hydrochloride at 5 mM (TCEP·HCl, Thermo Scientific Chemicals) at 37 °C for 20–30 min; newly exposed cysteines were, where indicated, re-alkylated with IAM at 5–10 mM for 10 min in the dark. Newly exposed cysteines were optionally capped with a second IAM addition (5–10 mM, 10 min). Urea was then diluted to ≤1 M with 50 mM ammonium bicarbonate, and sequencing-grade trypsin was added at a 1:50-1:100 enzyme: substrate ratio for 8–16 h at 37 °C. Digests were acidified to 0.5-1% formic acid, optionally desalted on C18 tips, dried, and reconstituted in 0.1% formic acid for LC–MS analysis.

Peptides were analyzed on an Agilent 6230 LC/TOF-MS operated in positive ion mode. Data were processed with Agilent MassHunter Qualitative Analysis using Find by Formula with a mass tolerance of ±10 ppm, targeting the WT tryptic peptides VFNDVR and DIIQR and the mutant peptide VFNDVCDIIQR carrying either IAM or BTFMA modifications. Assignments required agreement in monoisotopic mass, isotopic distribution, charge state (z = 2-3), and retention time, with BTFMA-modified peptides consistently eluting later than IAM-modified counterparts. Where available, MS/MS spectra were searched with fixed cysteine carbamidomethylation and a variable modification corresponding to the BTFMA mass shift, with peptide-spectrum matches filtered at 1% FDR and modification-site localization scored.

### Data Representation

All statistical tests, such as GTP hydrolysis assessments, were conducted using GraphPad Prism 9.0. The central point of all data points provides the mean value with standard deviation (s.d.) for all data, unless otherwise specified. Visualization of the atomic models (figures and videos) was made using PyMOL (The PyMOL Molecular Graphics System, Version 2.0, Schrödinger, LLC).

## Acknowledgements

This work is funded by the National Institutes of General Medical Sciences (1R01GM149659, L. Y.; R35GM128641, C.Z) and National Institute of Environmental Health Sciences (1R21ES035378, L. Y.) and start-up funding from the University of South Florida. We would also like to express our gratitude to Dr. Jinfa Ying at the National Institute of Diabetes and Digestive and Kidney Diseases for his advice on NMR acquisition, and to Dr. Heng Liu in the Department of Chemistry at the University of South Florida for his assistance with LC-MS data collection. Additionally, we thank Drs. Scott Prosser and Oliver Ernst from the University of Toronto, Canada, for their constructive comments on the manuscript.

## Author Contributions

L. Y. conceptualized and designed the study. L. Y. also processed some NMR and biochemical data. X.W. prepared and purified receptors and different Gα constructs. X. W. also performed NMR spectral acquisition, assisting with data processing, and spectral analyses in addition to biochemical assessments. W. S conducted GDP binding and nucleotide exchange experiments. N. H assisted in protein preparation and NMR data acquisitions. P. C.W. and C. Z. prepared Gβγ proteins. Y. Z. assisted with the setup and parameter optimization of the NMR pulse sequence. L. Y. supervised receptor and G protein purification, NMR acquisition, spectral analysis, and biochemical assays. L. Y. wrote the manuscript, and all authors contributed to the revisions.

## Competing Interests

The authors declare no competing financial interests.

## Notes

### Competing Interest Statement

The authors have declared no competing interest.

