## Supplementary material for "Ligand-Specified Signaling Efficacy Defined by Unique Transitions in G Protein Conformations": Supp

### Supplementary Materials

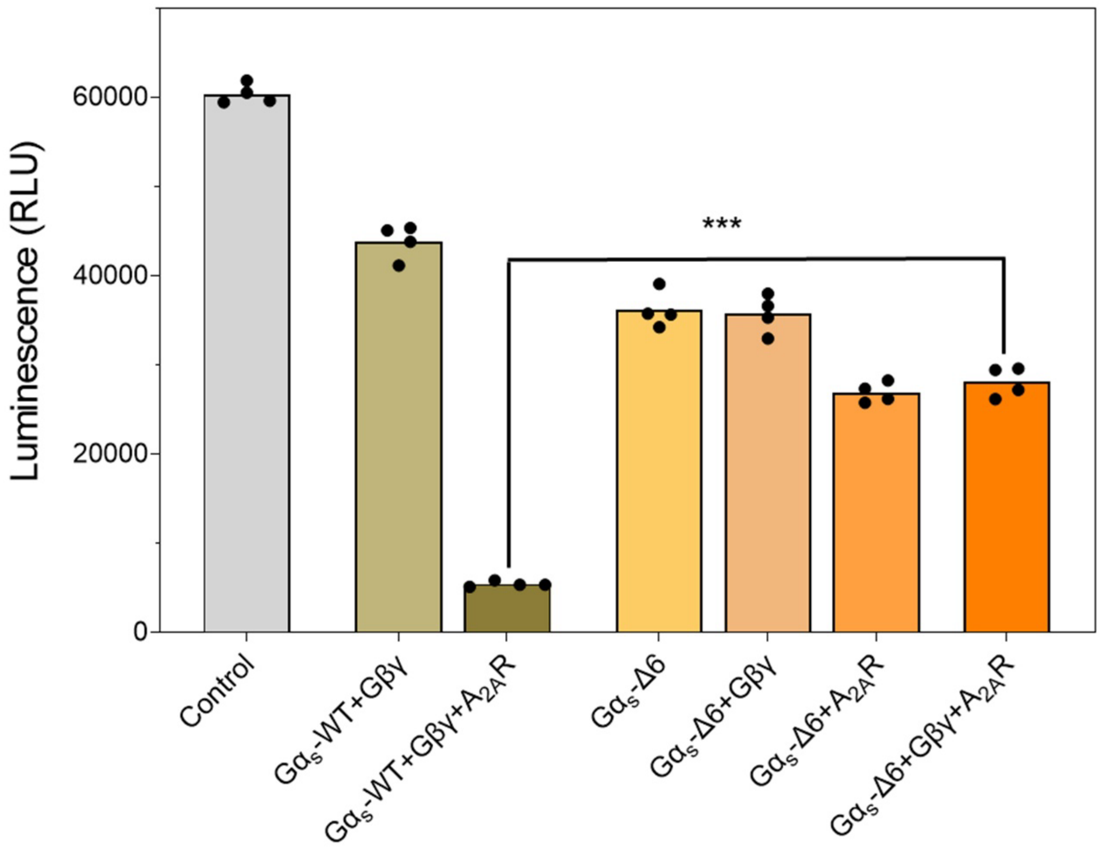


**Extended Data Fig.1. GTP hydrolysis assessment of cysteine-minimized Gα$_s$-Δ6, in comparison to the wild-type G protein.**

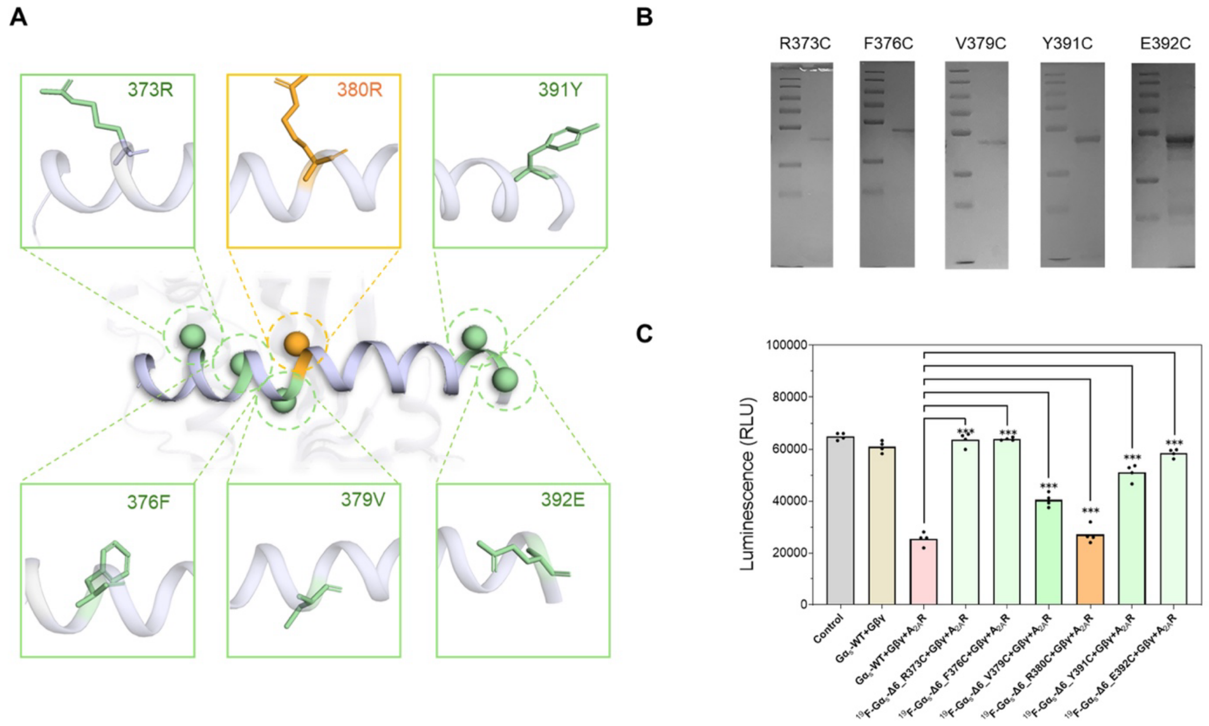


**Extended Data Fig.2. The G$\alpha_s$ construct screening for $^{19}$F-labeling. A.** Additional five sites were chosen to introduce single cysteine residues for BTFMA labeling. **B.** SDS-PAGE analysis of different $^{19}$F-labeled G$\alpha_s$ constructs. **C.** GTP hydrolysis assessments for various $^{19}$F-labeled G$\alpha_s$ constructs.

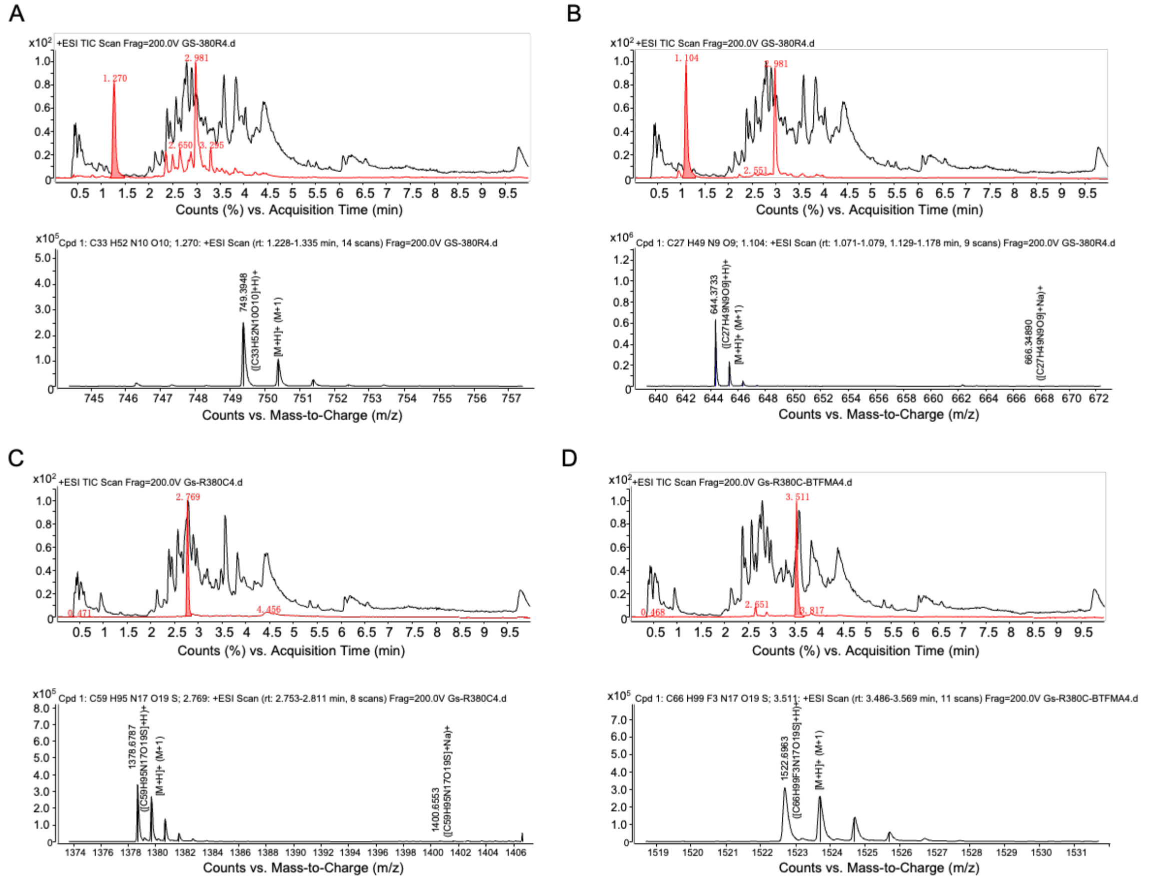


**Extended Data Fig.3. WT and BTFMA labeled $G\alpha_s$-$\Delta$6_R380C was assessed by LC-TOF mass spectroscopy. A.** WT (R380) short tryptic peptide VFNDVR: extracted-ion chromatogram (XIC; red, ±10 ppm) overlaid on total-ion chromatogram (TIC; black), with the corresponding MS spectrum at the XIC apex showing the charge-state envelope and monoisotopic peak. **B.** WT (R380) short tryptic peptide DIIQR, displayed as in **A**. These two panels establish the expected WT digestion products at the R380 cleavage site. **C.** Mutant R380C + IAM: detection of the longer peptide VFNDVCDIIQR carrying carbamidomethylation on Cys (+57.021 Da), with exact mass, isotope pattern, and co-eluting 2+ and 3+ charge states matching theory. **D.** Mutant R380C + BTFMA: detection of VFNDVCDIIQR carrying the probe adduct (+202.048 Da), eluting later than the IAM-modified analogue, consistent with increased hydrophobicity. Across panels, assignments required agreement in monoisotopic mass (≤10 ppm), isotope-pattern fit, and co-elution of z = 2 and z = 3 ions.

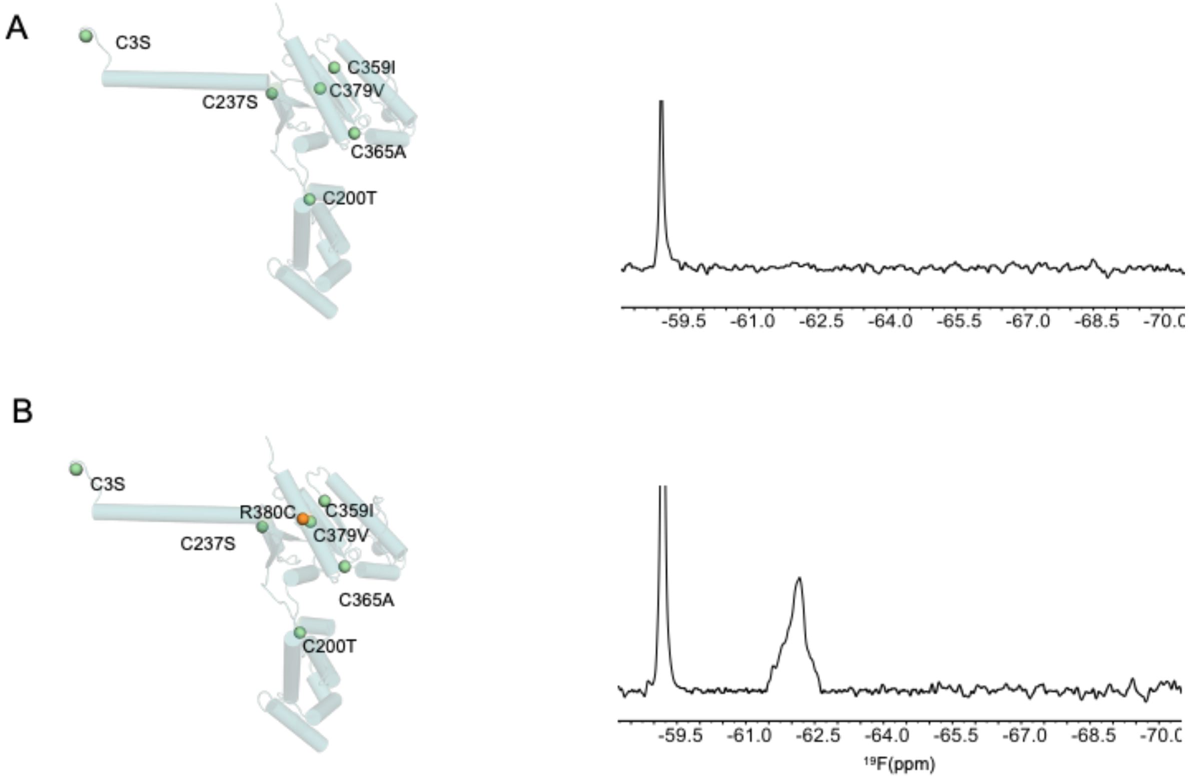


**Extended Data Fig.4. Non-cross labeling validation for the $G\alpha_s$-Δ6_R380C construct compared to $G\alpha_s$-Δ6. A.** Cysteine mutation sites of $G\alpha_s$-Δ6 construct and $^{19}$F-NMR after labeling with BTFMA. **B**. R380C labeling site based on $G\alpha_s$-Δ6 construct and $^{19}$F-NMR after labeling with BTFMA

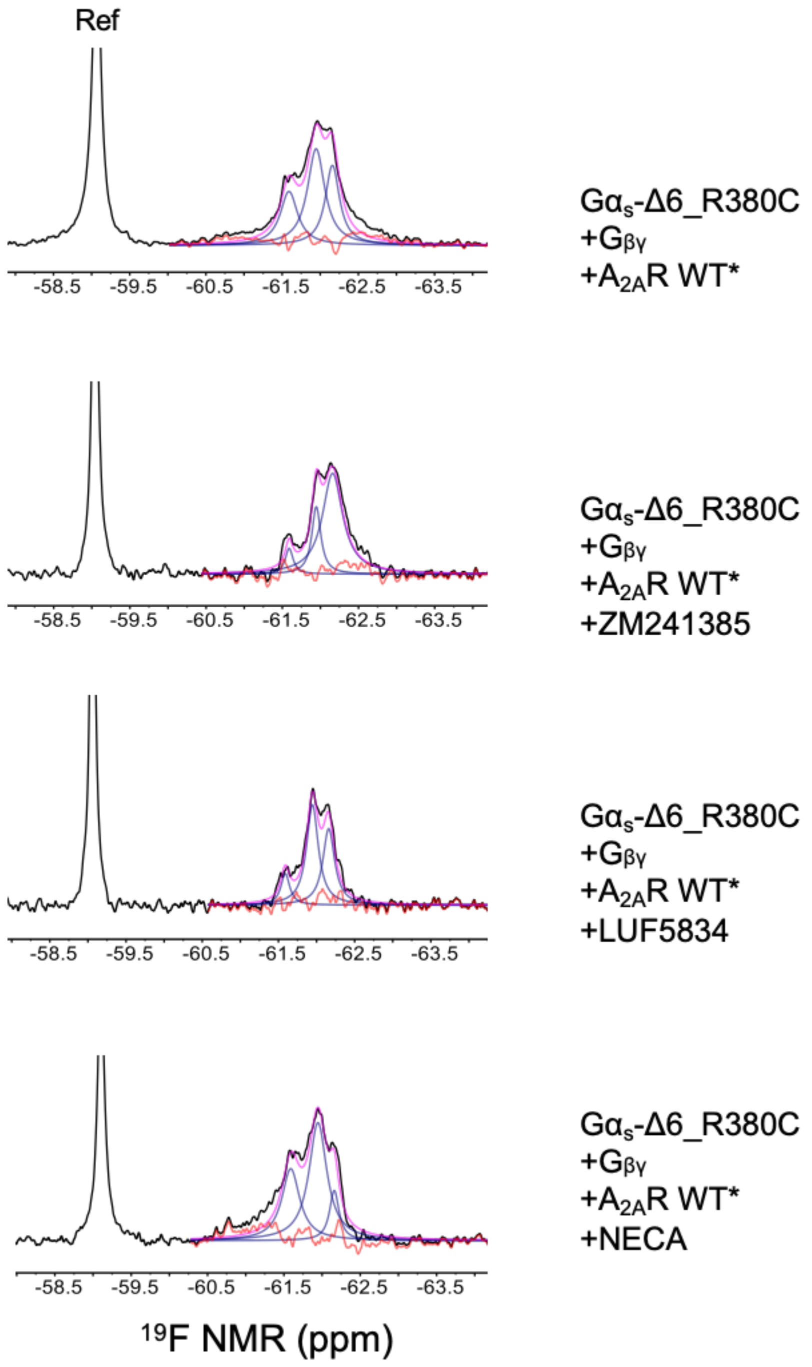


**Extended Data Fig.5. The $^{19}$F-G$\alpha_s$-Δ6_R380C NMR spectra of conformational transitions of the C$\alpha$5 helix in response to different ligand-bound $A_{2A}$R receptors.**

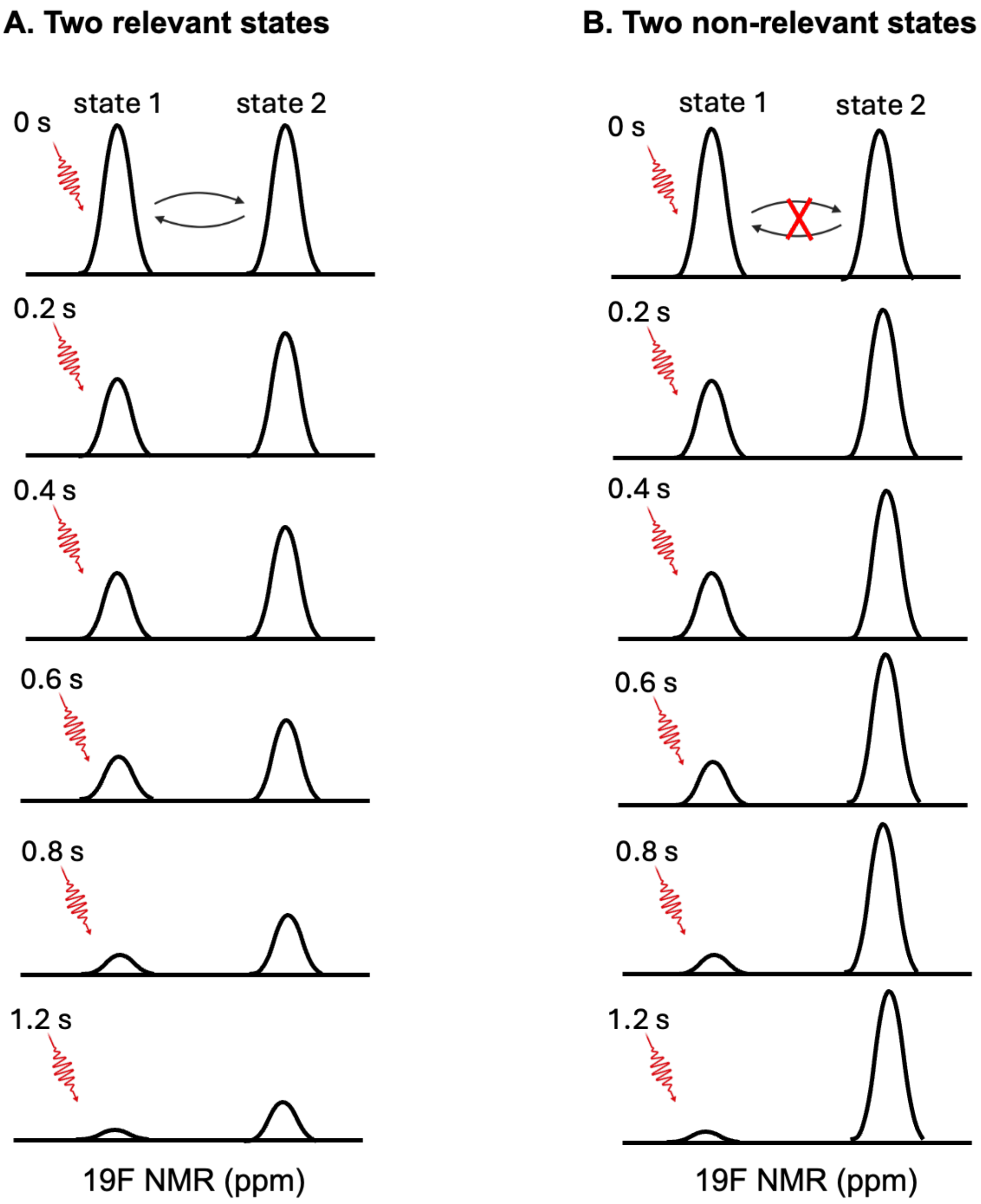


**Extended Data Fig.6. The working mechanism of 19F-CEST is demonstrated with two relevant and two non-relevant conformational states.**

### Extended Data Table 1. LC-TOF MS detection of R380/C380 labeling

| Sample | Peptide | Modification | Unmodified M (neutral) | Final M (neutral) | z=1 | z=2 | z=3 |
|---|---|---|---|---|---|---|---|
| WT | VFNDVR | none | 748.3868 | 748.3868 | 749.394 | 375.2007 | 250.4695 |
| WT | DIIQR | none | 643.3653 | 643.3653 | 644.373 | 322.6899 | 215.4624 |
| R380C + IAM | VFNDVCDIIQR | +57.021464 | 1320.6496 | 1377.6711 | 1378.678 | 689.8428 | 460.2310 |
| R380C + BTFMA | VFNDVCDIIQR | +201.048 | 1320.6496 | 1521.6976 | 1522.705 | 761.856 | 508.240 |

**Extended Data Table 2. Plasmids used in this study.**

| Plasmids | Source |
|---|---|
| pPIC9K_316_V229C | Ye, et al., Nature, 2016 |
| pBAD-MBP-TEV-G$\alpha_s$-Δ6-R380C | This study |
| pBAD-MBP-TEV-G$\alpha_s$-Δ6-R373C | This study |
| pBAD-MBP-TEV-G$\alpha_s$-Δ6-F376C | This study |
| pBAD-MBP-TEV-G$\alpha_s$-Δ6-V379C | This study |
| pBAD-MBP-TEV-G$\alpha_s$-Δ6-Y391C | This study |
| pBAD-MBP-TEV-G$\alpha_s$-Δ6-E392C | This study |
| pBAD-MBP-TEV-G$\alpha_s$-Δ6-WT | This study |

### Extended Data Table 3. Primer sequences used in this study

| Primers | Sequences (5'-3') |
|---|---|
| Gα-F-NcoI | TGGGCTAAGGAGGAATTAACCATGGGTCACCAT |
| Gαs-R-KpnI | CGGGGTACCTTATTACAGCAGTTCGTACTGACGCAGGTGCAT |
| pBAD-MBP-TEV-$G\alpha_s$-C3S | CTTCCAGGGTGCTATGGGTAGCCTGGGTAA |
| pBAD-MBP-TEV-$G\alpha_s$-C200T | CGTCTGACCAGGACTTACTGCGTACCCGTGTTCTGAC |
| pBAD-MBP-TEV-$G\alpha_s$-C237S | GGTAACGTCGTTGAAGCTCTGGATCCATTTACGAC |
| pBAD-MBP-TEV-$G\alpha_s$-C359I | TGACGGTCGTCACTACATCTACCCGCACTTCACC |
| pBAD-MBP-TEV-$G\alpha_s$-C365A | ACCCGCACTTCACCGCCGCTGTTGACACCG |
| pBAD-MBP-TEV-$G\alpha_s$-C379V-R380C | TCCGTCGTGTTTTCAACGACGTCTGTGACATCATCCAGCGTATG |
| pBAD-MBP-TEV-$G\alpha_s$-C379V | CGTCGTGTTTTCAACGACGTCCGTGACATCATCCAGCG |
| pBAD-MBP-TEV-$G\alpha_s$-R373C | CGTCGTTGAAAACACGACAGATGTTTTCGGTGTCAAC |
| pBAD-MBP-TEV-$G\alpha_s$-F376C | CAGACGTCGTTGCAAACACGACGGATGTTTTCGG |
| pBAD-MBP-TEV-$G\alpha_s$-V379C | CGCTGGATGATGTCACAGCAGTCGTTGAAAACACGACG |
| pBAD-MBP-TEV-$G\alpha_s$-Y391C | CGTATGCACCTGCGTCAGTGCGAACTGCTG |
| pBAD-MBP-TEV-$G\alpha_s$-E392C | CGTATGCACCTGCGTCAGTACTGCCTGCTGTAATAAGGTACCATA |
